# Dietary supplementation with products of Citrus reticulata “Chachi” for improving the fecal microbiome of weaned piglets

**DOI:** 10.1101/2020.04.14.040881

**Authors:** Huafeng Lin, Gang Li, Haizhen Wang, Lili Li, Xun Chen, Lei Shi, Yong Tang, Yanlei Chang, Jie Yang, Yuqi Liu, Hanyue Gong

**Affiliations:** Department of Biotechnology, College of Life Science and Technology, Jinan University, Guangzhou, Guangdong 510632, PR China; Institute of Food Safety and Nutrition, Jinan University, Guangzhou 510632, PR China; Institute of Biomedicine and Department of Cell Biology, Jinan University, Guangzhou 510632, P.R. China; Food and Health Engineering Research Center of the State Education Ministry, School of Life Science, Sun Yat-Sen University, Guangzhou, PR China; Key Laboratory of Functional Protein Research of Guangdong Higher Education Institutes, Institute of Life and Health Engineering, College of Life Science and Technology, Jinan University, Guangzhou 510632, PR China

## Abstract

Nutritional interventions play a critical role in modifying the intestinal microbiome of host animals. This study was conducted to interrogate the physiological effects on fecal microflora of weaned piglets via the dietary supplemented with two types of products of Citrus reticulata “Chachi”, respectively. For this purpose, A total of 72 piglets with uniform sizes were randomly assigned to four dietary treatment groups consisted of a negative control group (NCG), a fermented citrus reticulata “Chachi” pulp group (FCRPG), a Citri Reticulatae Pericarpium group (CRPG) and a positive control group (PCG) in a 21-day feeding trial. After the raising experiment, fresh feces of piglets were analyzed systematically using multi-omics technologies. Metagenomics method with high-throughput compositional characterization indicated that the architecture and diversity of fecal microbiome were both influenced by these two additives and compound antibiotics. Metabolite analysis showed that FCRPG have an significant effects on fecal short-chain fatty acids (SCFAs) among four treatment groups. Results of functional proteomics approaches found that FCRPG presented the highest butyrate metabolic level, and CRPG showed the highest flavone and flavonol biosynthesis level in feces. In addition, NCP produced an effective effect on adjusting fecal microbiota profile. Consequently, our findings demonstrate that dietary supplementation with FCRP or CRP modulates the microbial taxa, metabolic and proteomic alterations in fecal microbiota of weaned piglets for health maintaining.

## Introduction

Swine is an important economic specie in livestock production worldwidely. The microbiome within digestive tract of each swine is a complex and miraculous system [1], which are usually associated with physiological functions such as energy metabolism, immune regulation and gene expression. The vast majority of commensal micro-bacteria are well adapted to the host environment but also can be shaped by various factors such as nutrient import and external circumstances [2, 3]. Weanling period is a special developmental stage throughout the entire lifecycle of pig, during which piglets have exhibited rapid growth, high demand for nutrition, fast energy harvesting capability [4], and succession of gut microbial composition [5-7]. However, common diseases such as digestive disorders and intestinal inflammation, often plague the piglet farming during this rearing phase. Gut microbiota obviously play the function of facilitating adaption of weanling piglets to fibrous ingredients and reducing the risk of colonization of enteric pathogens after weaning [8]. Actually, although antibiotics have been pervasively applied many decades for antimicrobial application and growth promoting for livetocks in China, European Union has banned their prophylactic use in modern swine industry for the potential public hazard of bacterial resistances and drug residues since 2006 [9, 10]. Thus, nutritional strategies have been employed as an efficient way to improve intestinal microbiota architecture for the purpose of restricting the use of antibiotics and enhancing the health of piglets except for alternative methods such as management and housing [11-13].

To our knowledge, fresh citrus fruits are often used directly in piggeries in tropical countries or regions such as Vietnam, Colombia, Guadeloupe, United States, Netherlands, but their quantity of usage is limited [14]. So far, citrus pulp are mainly utilized as energy source for ruminant livestock (such as cattle, sheep, etc.), and also as a dietary supplemention in pigs [15]. However, the usage method (dried citrus pulp vs. ensiled citrus pulp) and quantity of citrus pulp are well controlled and vary with the differences of mammalian digestive systems. For instance, Cerisuelo et al. (2010) reported that utilization of ensiled citrus pulp in the diets of growing pigs has no deleterious effects on the growth performance and meat quality, and shows potential benefits for gastrointestinal (GI) tract microorganisms [16].

According to the publications of the Committee on Herbal Medicinal Products (HMPC) at the European Medicines Agency (EMA), many medicinal plants have been conventionally used for the remedy of GI diseases [17]. Similarly in China, Citri Reticulatae Pericarpium (CRP), made from peels of Citrus reticulata “Chachi”, is one of the most famous Chinese medicines officially presented in the Chinese Pharmacopoeia [18]. Traditionally, CRP is used to ameliorate digestion metabolism and deal with certain respiratory disorders such as cough and phlegm [19]. Previous study revealed that CRP possesses approximately 140 chemical compounds such as flavonoids, essential oils, polysaccharides, carotenoids, vitamins, minerals and so on [20]. Recently, CRP has been developed for functional food additives as they possess many vital pharmacological properties such as anti-oxidative activity, anti-inflammatory activity, anticancer effect, anti-asthmatic activity and so on [20–22].

Fermented citrus reticulata “Chachi” pulp (FCRP), a type of by-products in the production of CRP, have been naturally fermented/ensiled for appropriate 15 days after being crushed and deseeded. Nutritional studies suggested that citrus pulp is a good source of sugars [23], flavanones [24], phenolic compounds [25], vitamins [26] and minerals [27], etc. In the pig production, FCRP can also be used as a dietary acidifier to minimise detrimental effects due to its high contents of antimicrobial organic acids [28]. In addition, the fermentation technique is a kind of good way to reduce anti-nutrients (e.g., tannin and phytate [29]) in citrus reticulata “Chachi” pulp, and to enhance the levels of beneficial microorganisms (e.g., lactic acid) slightly so as to improve diet palatability and digestibility.

Nowadays, with the advance of multi-omics technologies, we are able to intensively dissect the correlations among fecal microbiota composition, fecal proteomics profile and microbial metabolites, which in relation to the disease and health of weaned piglets.

The objective of present study is to assess effectiveness and functionality by the dietary inclusion of processed products of Citrus reticulata “Chachi” for optimizing fecal bacterial community composition of weaned piglets. For this purpose, we integrated advanced multi-omics technologies to evaluate the influences of dietary administration of FCRP, CRP and antibiotics, respectively, to reveal the relationship between dietary intake components and fecal microbial composition. Finally, various functional proteins were identified by searching proteomics databases, revealing the presence of several functional pathways that linked these exogenous additives (FCRP, CRP and antibiotics) to fecal microbiota.

## Materials and methods

### Animal experiments

The animal care and procedures used in this investigation were performed in term of the guidelines of the China Animal Protection Association, and were approved by the Institutional Animal Ethics Committee of Jinan University.

### Animal raising and operations

Under uniform conditions of husbandry, 72 healthy piglets (Duroc × Landrace × Large White), which weaned at the age of 3 weeks with mean body weight of 6.38 ± 0.18 kg, were stochastically allocated to four treatment groups, with each group comprising three replicated pens of 6 piglets. These castrate piglets were reared in cement floor pens (2.0 m × 2.5 m) with ad libitum access to clean drinking water for 21 days. Prior to the trial, all piglets were received to the basal diets formulated in accordance with recommendations of NRC (2012) [30]. During the experiment, ambient air temperature in the open pens was maintained at 20–26 °C. The routine work of hog lots are implemented according to the pigsty management procedures.

### Ingredients preparation and formulation design

FCRP and CRP used in the present trial were supplied in part by Xinhui “Yi Pintang” tea ceremony factory (Jiangmen, China) and Xinhui “Gan Cheng” tea ceremony factory (Jiangmen, China). The Citrus reticulata “Chachi” fleshes were crushed, deseeded and stored in a polythene food barrel with a bottom diameter of 1.0 m and height of 2.0 m for continuous fermentation of about half a month. CRP that stored hermetically and aged for over 3 years were smashed into powder prior to use.

Four dietary formulations fed for piglets (Table 1) were set as follows: negative control group (NCG), fermented citrus reticulata “Chachi” pulp group (FCRPG), Citri Reticulatae Pericarpium group (CRPG) and positive control group (PCG).

**Table 1.** Composition and nutrient levels of experiment diet (air-dry basis).

| Ingredients | Diets |  |  |  |
| --- | --- | --- | --- | --- |
|  | NCG | FCRPG | CRPG | <sup>4</sup>PCG |
| Corn | 56.0 | 51.5 | 51.0 | 56.0 |
| SM(45% CP) | 26.0 | 26.0 | 26.0 | 26.0 |
| Wheat bran | 8.0 | 8.0 | 8.0 | 8.0 |
| Rice bran | 6.0 | 6.0 | 6.0 | 6.0 |
| FCRP | - | 4.5 | - | - |
| CRP | - | - | 5 | - |
| <sup>1</sup>Vitamin premix | 0.5 | 0.5 | 0.5 | 0.5 |
| <sup>2</sup>Mineral premix | 0.5 | 0.5 | 0.5 | 0.5 |
| Antibiotics | - | - | - | + |
| <sup>3</sup>Others | 3.0 | 3.0 | 3.0 | 3.0 |
| Total | 100 | 100 | 100 | 100 |
| Nutritional composition (NC) |  |  |  |  |
| Digestible energy<br>(DE, KJ/kg) | 14.98 | 14.65 | 14.93 | 14.96 |
| Crude protein (CP, %) | 18.46 | 18.11 | 18.43 | 18.45 |
| Ether extract (EE, %) | 4.25 | 4.02 | 4.11 | 4.23 |
| Ash (%) | 3.81 | 3.90 | 3.96 | 3.80 |

<sup>1</sup>Vitamin premix (mg/Kg diet): VA, 8000 IU; VD3, 1000 IU; VE, 60 IU; VK3, 0.7; VB1, 1.1; VB2, 2.3; VB6, 1.7; VB12, 0.02; VB5, 21; Niacin, 10;
<sup>2</sup>Mineral premix (mg/Kg diet): Mg, 160; Cu, 11; Co, 0.45; Zn, 100; Mn, 30; Fe, 145; Se, 0.1; I, 0.6.
<sup>3</sup>Others (mg/Kg diet): lysine: 1.3; methionine: 0.08; threonine: 0.08; Ca(H<sub>2</sub>PO<sub>4</sub>)<sub>2</sub>: 20.3; CaCO<sub>3</sub>: 3; NaCl: 3; FeSO<sub>4</sub> • 4H<sub>2</sub>O: 0.24; TiO<sub>2</sub> (IV): 2.0.
<sup>4</sup>PCG: refers to adding 0.11% compound antibiotics on the basis of NCG diet, and the components of this antibiotics are: bacitracin zinc / polymyxin sulfate =5/1.

### Sample collection and processing

Flesh stool specimens were immediately collected either using a sterilized cryopreservation tube with a collection brush during defecation or via rectal massage method noninvasively [31]. Three samples from different piglets of same treatment group were merge into one tube, which was quickly snap-frozen in liquid nitrogen and transported to laboratory for DNA extraction, the remaining samples are stored at –80 °C for further processing.

### DNA extraction and 16S rRNA gene sequencing

Bacterial DNA was extracted from unprocessed piglets’ feces using QIAamp DNA Mini Kit (Qiagen, Hilden, Germany) following the internal quality SOP of products. The final DNA concentrations were determined using a Nanodrop 2000 UV/Vis spectrophotometer (Thermo Fisher Scientific, Wilmington, United States) on the basis of the absorbance of 260 nm to 280 nm, and agarose gel electrophoresis test was used for evaluation of the DNA integrity as previously described [32]. Negative controls were performed for the extraction process and evaluated by gel analysis after polymerase chain reaction (PCR) amplification. For deep sequencing analysis, the V3–V4 hypervariable regions of the 16S rRNA gene was amplified using forward (338F, 5’-ACTCCTACGGGAGGCAGCAG-3’) and reverse (806R, 5’-GGACTACHVGGGTWTCTAAT-3’) primers conducted on a QuantStudio™ 6 Flex (Life Technologies, U.S.) PCR system. PCR reactions were performed in triplicate 20 µL mixed solution containing 10 ng template DNA, 4 µL 5 × FastPfu buffer, 2 µL dNTPs (2.5 mmol/L), 0.8 µL each primer (5.0 µmol/L), 0.4 µL FastPfu polymerase and ddH_2_O. The reaction conditions for PCR were 95 °C for 3 min for an initial denaturation, followed by 27 cycles of denaturation at 95 °C for 30 s, primer annealing at 55°C for 30 s, extension at 72 °C for 45 s, and a final elongation for 10 min at 72°C. The PCR products were extracted from 2% agarose gels, and by following the manufacturer’s instructions, the further purification and quantification were used the AxyPrep DNA Gel Extraction Kit (Axygen Biosciences, Union City, CA, U.S.) and QuantiFluor™–ST (Promega BioSciences LLC, Sunnyvale, CA, U.S.), respectively. In the PCR reactions, samples without template and those contained known 16S rRNA gene sequences were respectively used as negative and positive controls. DNA was preserved in a –80°C refrigerator for the downstream treatments.

### SCFAs quantification

Fecal samples of piglets were pretreated (three sample repetitions for each treatment) according to the method of zijlstra and colleagues [33]. Using 4-methylisovaleric acid as an internal standard, the concentrations of fecal SCFAs were determined by gas chromatography.

### Total protein extraction and mass spectrometry analysis

Approximately 0.3g fecal samples of each group are used for the extraction of total protein according to the method of haange and co-authors [34]. The protein extracts were separated and purified using sodium dodecyl sulfate-polyacrylamide gel electrophoresis (SDS-PAGE), and then the single band cutted in gel state was digested by trypsin [35, 36]. The desalted peptide assortments were analyzed by the application of label-free proteomic quantitative techniques on a mass spectrometry-based proteomic research platform (Thermo Scientific™ Orbitrap Fusion™).

### Mass spectrometry data analysis

The mass spectrum data were extracted by protein search software (Mascot 2.3.02), and then the peptides/proteins data were under quality controlled, and identified with database searching (NCBI RefSeq library and UniProtKB Library) by using Scaffold Software. Applied “false discovery rate (FDR) < 1.0%” as the screening standard for target peptides via local FDR algorithm of Scaffold. The intensities obtained from the experiment were normalized within all aquisition runs. Use “fold change ≥ 1.5 and *P* < 0.05” as the standard of screening differential proteins.

### Bioinformatics analysis

All quantified peptides were subjected to Unipept 4.0 for taxonomic assignment based on the principle of Lowest Common Ancestor (LCA) [37]. Cluster of Orthologous Groups of proteins (COGs) is used to functionally classify different bacterio-proteins. The abundance of each category consisted of the sum of the protein intensities of all COGs in the category. Ultimately, the identified COGs were applied to the Kyoto Encyclopedia of Genes and Genomes (KEGG) metabolic pathway website (https://www.kegg.jp/) for downstream analysis.

### Statistical analysis

Data were analyzed via one-way ANOVA using SPSS 19.0 software (IBM Corp., New York, NY, United States). Data are presented as means ± S.D., and differences were regarded as statistically significance at *P* < 0.05. Venn diagram analysis was performed to present unique and shared OTUs in the fecal microbiome among four treatment groups. Heatmap was generated with the R-package gplots at the genus level.

## Results

### Microbiota composition and diversity

After the raw paired-end reads were quality filtered and assembled, 458,016 effective sequences were obtained from 12 piglet’s fecal specimens. A mean of 38,168 reads per sample were acquired with iSeq 100 (illumina) after sequencing the V3-V4 region of the 16S rRNA gene from fresh feces. A total of 573 OTUs (Fig. 1) which were grouped in 18 Phyla, 43 Classes, 105 Orders, 180 Families and 410 Genera.

**Fig. 1.**
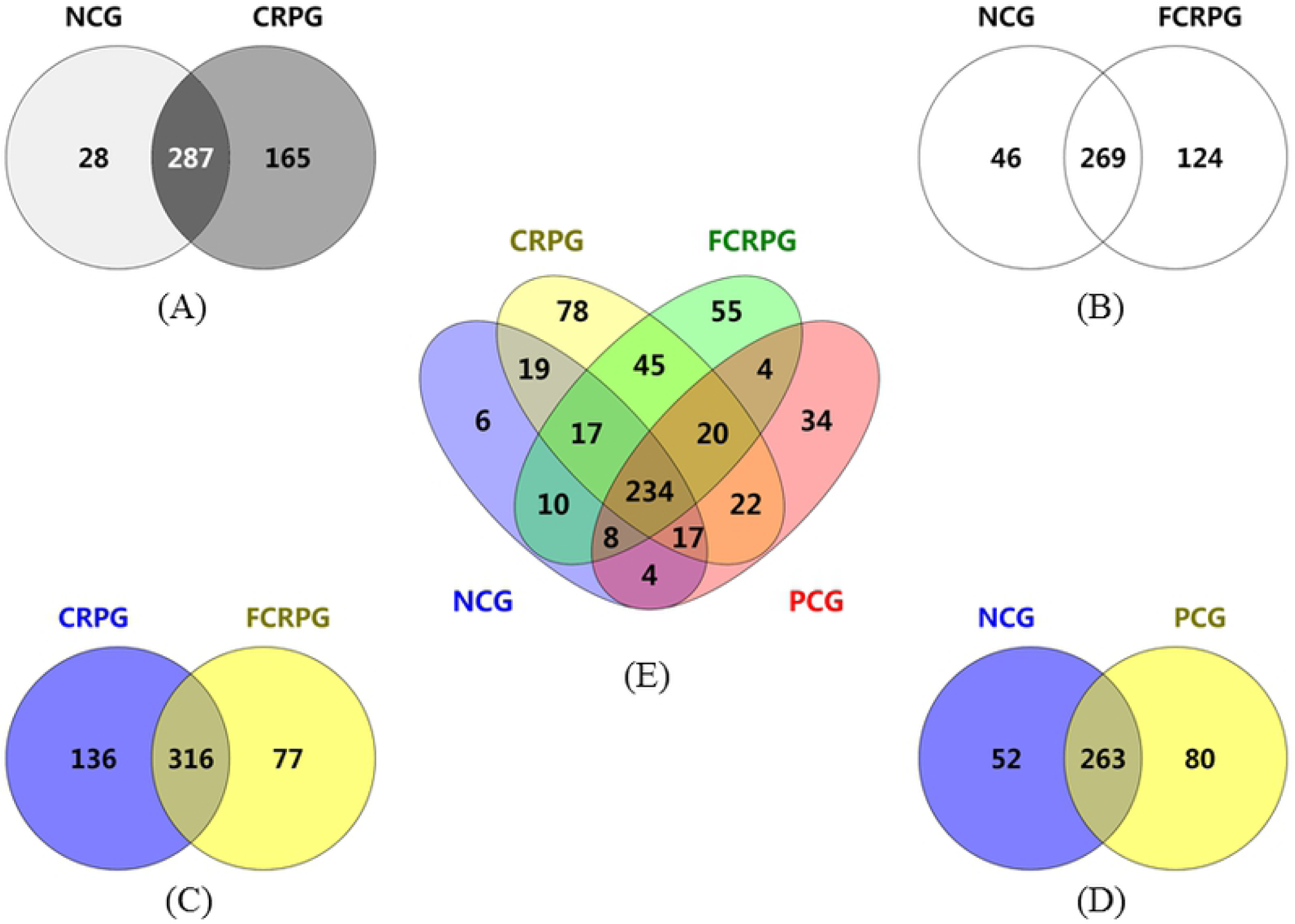
The venn diagrams showing the unique and shared OTUs among the four treatment groups. (A) Between NCG and CRPG, (B) Between NCG and FCRPG, (C) Between FCRPG and CRPG, (D) Between NCG and PCG, (E) Among four treatment groups.

### Bacterial alpha diversity analysis

The coverage of sequencing in the samples is over 99.9%. Table 2 showed the diversity indexes of fecal microorganisms among four dietary treatment groups (NCG, FCRPG, CRPG, and PCG).

**Table 2.**
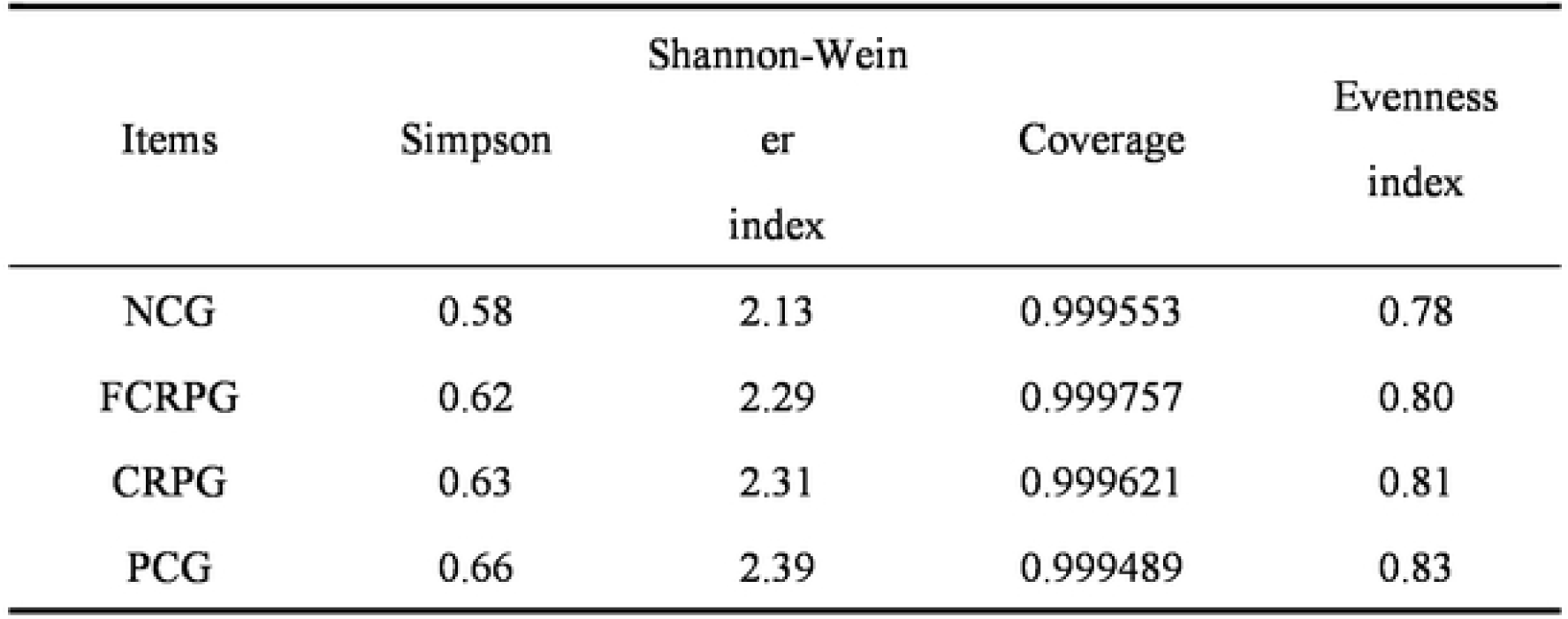
Diversity indexes of fecal microbiota for weaned piglets.

| Items | Simpson | Shannon-Weiner index | Coverage | Evenness index |
| --- | --- | --- | --- | --- |
| NCG | 0.58 | 2.13 | 0.999553 | 0.78 |
| FCRPG | 0.62 | 2.29 | 0.999757 | 0.80 |
| CRPG | 0.63 | 2.31 | 0.999621 | 0.81 |
| PCG | 0.66 | 2.39 | 0.999489 | 0.83 |

The Simpson diversity indices of NCG was obtained a lower value than three additional groups (FCRPG, CRPG, and PCG). According to the formula of Simpson’s diversity index: 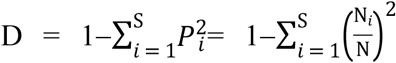, the Simpson diversity indices (represented with the letter ‘D’) is positively correlated to species amount (represented with the letter ‘S’) and species evenness index. Therefore, NCG had the lowest values of species amount and species evenness compared to three additional treatment groups. Similarly, the Shannon-Weiner index primarily reflects the species diversity of bacterial community. The relative lowest value of Shannon-Weiner index acquired in NCG indicated that FCRPG, CRPG and PCG had higher bacterial diversity index values, which illustrating FCRP, CRP and compound antibiotics are capable of improving the bio-diversity of intestinal microorganisms to a certain degree.

Overall, there is no significant difference measured in fecal microbial diversity indexes among dietary treatment groups (NCG, FCRPG, CRPG, and NCG) (*P* > 0.05). However, these data still indicated that the inclusions of FCRP, CRP and compound antibiotics to the diet all have modulated the fecal microorganisms of weaned piglets, resulting in modifications of fecal microbial diversity indexes, which in turn affects the health growth of weaned piglets.

### Venn diagram analysis

The Venn Diagram showed that the unique and shared OTUs of fecal microbiota for weaning piglets among four treatment groups (NCG, FCRPG, CRPG and PCG). Obviously, significant overlap patterns were observed between any two groups (NCG and FCRPG, NCG and CRPG, FCRPG and CRPG, NCG and PCG) (Fig. 1). Among which, 269 OUTs are shared by NCG and FCRPG (Fig. 1B), 287 OUTs are shared by NCG and CRPG (Fig. 1A), 316 OUTs are shared by FCRPG and CRPG (Fig. 1C), while 263 OUTs are shared by NCG and PCG (Fig. 1D). This illustrated that the majority of common OTUs (234 out of 573) may probably be the permanent residents of rectum of weaned piglets (Fig. 1E). Among 573 OUTs, NCG owned only 6 unique OUTs, while FCRPG, CRPG and PCG had 55, 78 and 34 unique OUTs, respectively. These indicated that FCRPG, CRPG and PCG increased their respective fecal microbiota diversities in varying degrees compared with NCG. Specially, some parts of bacterial OUTs derived from dietary supplemention of FCRP or CRP may pass through the whole intestinal tract of piglets, and thus eventually appear in their feces, as a consequence of the digestive system and immune system of weaning piglets are relatively functionally imperfect. In addition, there are also evidences showed that the intervention of compound antibiotics can disturb the intestinal microbiota composition with numerous possible outcomes in swines [38, 39]. Here in our experiment, a slight increase of fecal microbial abundance was observed in weaned piglets which fed diet contained a certain amount of antibiotics.

### Thermal imagine analysis

Some certain indigenous bacterial genera of feces were correlated with altered dietary ingredients. As shown in Fig. 2, heatmap analyses of the relative abundance of fecal microorganisms were displayed at the genus level. Among four treatment groups, the genus Prevotella presents similar high abundance (bright green band), while the genus Bilophila (especially Bilophila wadsworthia), which has been reported to trigger inflammatory bowel disease in mouse models [40], exibits similar low abundance (red band).

**Fig. 2.**
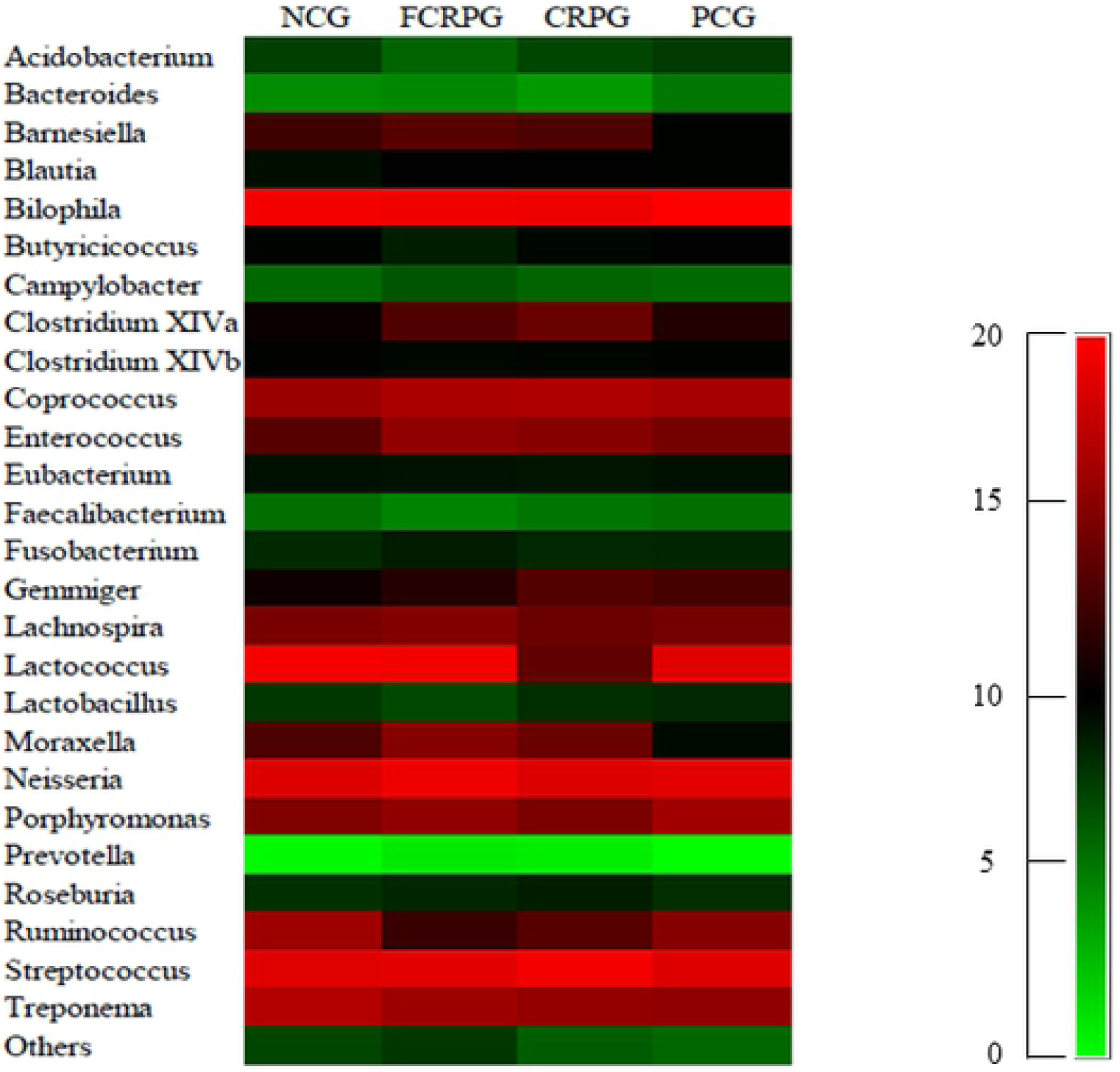
Genus heatmap analyses showing the fecal microbiota memberships in piglets’ feces. The heatmap plot depicts the relative percentage (%) of each fecal bacterial genera (vertical-axis clustering) within each treatment group (horizon-axis clustering). The color of the spots in the right panel represents the relative abundance values (%) of the dominant genera in corresponding treatment group.

### Taxonomic classification of fecal bacteria

As shown in Fig. 3, the effects on fecal microorganisms of weaned piglets of four dietary groups (NCG, FCRPG, CRPG, and PCG) at the genus (Fig. 3c), family (Fig. 3b) and phylum (Fig. 3a) levels. From the horizontal point of view, the stacked bar plots clearly showed the relative percentage contents of fecal microbes at each genus, family and phylum levels. According to the sequencing data, the Lactobacillus, Lactococcus, Prevotella, Acidobacterium, Streptococcus, Micrococcus of four treatment groups were compared longitudinally (Fig. 4). Collectively, FCRP has obviously increased the relative abundance of Lactobacillus and Acidobacterium in piglets’ feces, and decreased the relative abundance of Streptococcus, while CRP has obviously increased the relative percentage of Lactococcus and Prevotella. In the present trial, these two dietary addititves (FCRP and CRP) are particularly associated with changes in the number of Lactobacillus, Lactococcus, Micrococcus and Streptococcus in the feces of weaned piglets. In addition, the compound antibiotics can also disturb the microbial memberships in the feces of piglets. Consequently, there is a certain corresponding relationship between specific dietary components and specific types of microbial flora.

**Fig. 3.**
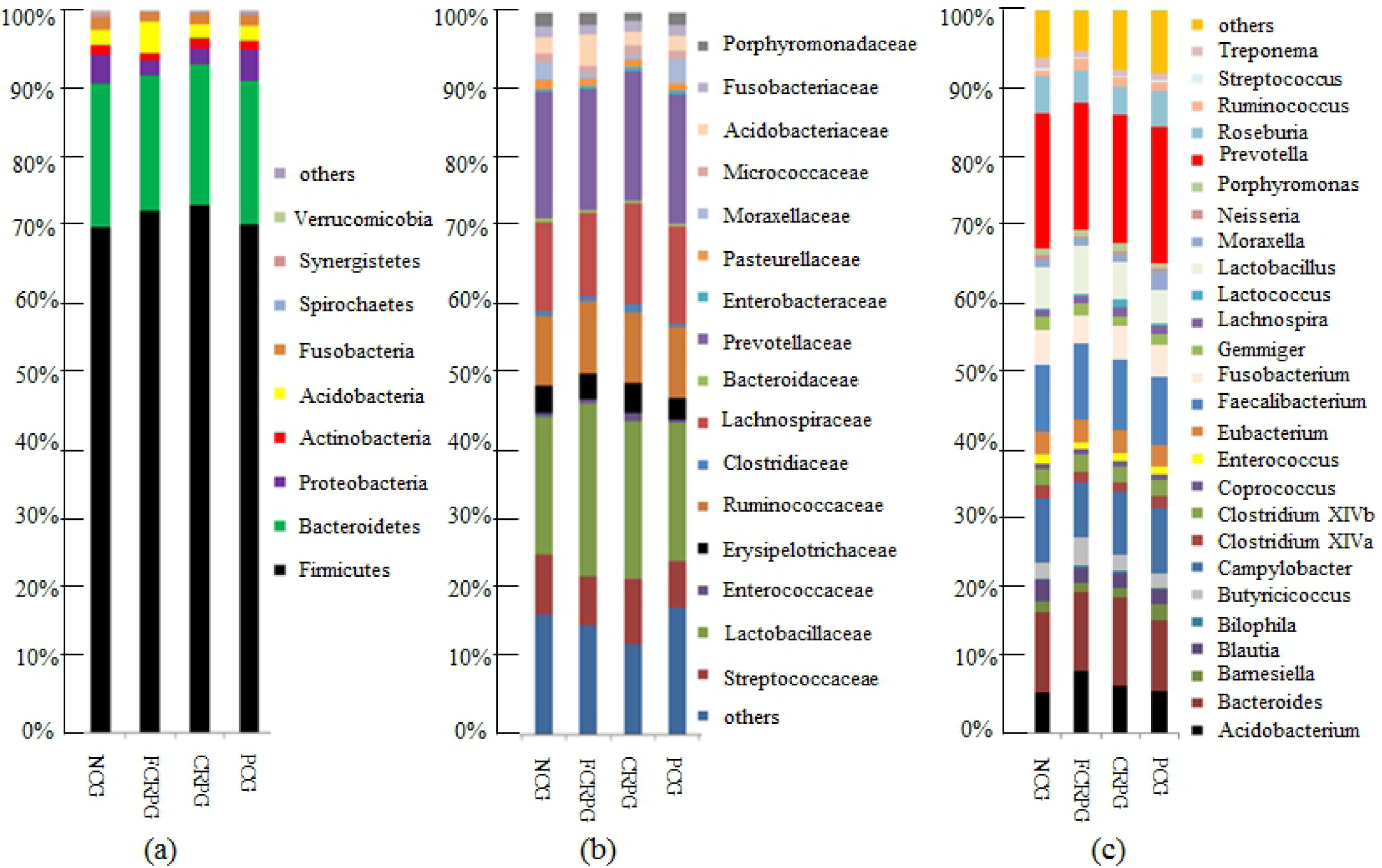
Bar-plot analysis showing fecal microbial community structure of weaned piglets among four treatment groups at the Phylum, Family and Genus levels. (a) Barplot at the phylum-level. (b) Barplot at the family-level. (c) Barplot at the genus-level. Each bar represents the average relative abundance (%) of each bacterial taxon within per treatment group.

**Fig. 4.**
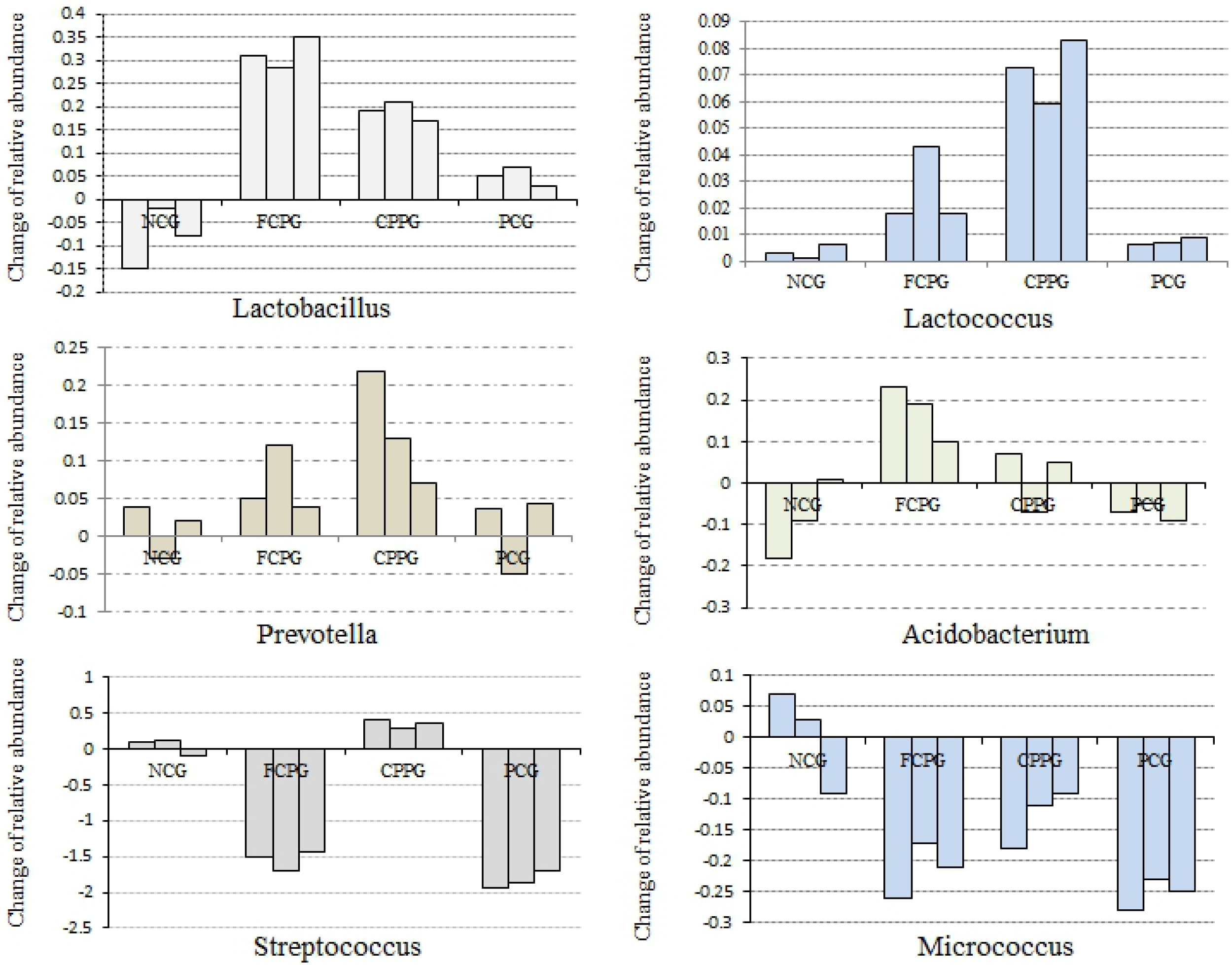
Influences on the relative abundance of microbiota among dietary treatment groups. The bar charts shows bacterial taxa that were affected by the dietary supplement of FCRP or CRP. Bars illustrating the change in relative abundance between samples collected at day 0 and day 21 for Lactobacillus, Lactococcus, Prevotella, Acidobacterium, Streptococcus and Micrococcus among four treatment groups.

### Analysis of the fecal microbial metabolites in weaned piglets

The fecal SCFAs profiles in weaned piglets is given in Table 3, which presenting the concentration of total SCFAs and its components in feces. There were significant differences (*P* < 0.05) in the concentrations of acetic acid, propionic acid, butyric acid and total SCFA between FCRPG and three additional groups. Significant differences (*P* < 0.05) also found in the contents of acetic acid and total SCFAs between CRPG and NCP. However, no significant differences (*P* > 0.05) of propionic acid and butyric acid between CRPG and NCP.

**Table 3.** The SCFAs concentration in piglets’ feces among dietary treatment groups.

| Categories | NCG | FCRPG | CRPG | PCG | *P*-Values |
| --- | --- | --- | --- | --- | --- |
| Acetic acid | 249.76±2.55<sup>a</sup> | 281.70±2.50<sup>c</sup> | 266.80±2.76<sup>b</sup> | 247.97±2.70<sup>a</sup> | 0.000 |
| Propionic acid | 94.97±3.91<sup>a</sup> | 107.93±5.68<sup>b</sup> | 98.80±2.19<sup>a</sup> | 98.40±0.75<sup>a</sup> | 0.013 |
| Butyric acid | 55.8±1.3<sup>a</sup> | 61.9±1.10<sup>b</sup> | 57.63±1.15<sup>a</sup> | 56.30±0.98<sup>a</sup> | 0.001 |
| Total SCFAs | 400.53±4.58<sup>a</sup> | 451.53±7.60<sup>b</sup> | 423.23±6.05<sup>c</sup> | 402.67±1.32<sup>a</sup> | 0.000 |

Note: Three duplicates for each treatment group.

### Mass spectrometry analysis of fecal proteomics

In this study, 2540 non-redundant proteins were identified, including 2,465 bacterial proteins (97.05%) and 75 pigling’s proteins (2.95%). Proteins/peptides having two or more peptide segments are artificially classified as “the effective proteins”. Based on such principle, 1,871 proteins were identified, 1812 of which were from bacteria (96.85%), and 59 from piglets (3.23%), excluding possible contaminants. Using “fold change ≥ 1.5 and *P* < 0.05” as the standard of filtering differential proteins, 241 differently expressed proteins in total were identified, of which 230 (95.43%) are from fecal bacteria and 11 (4.56%) are from piglets.

In addition, a total of 5,106 peptides were identified, of which 4,850 were bacterial peptide segments. Peptides that belonging to the 230 differential bacterial proteins were artificially divided into significantly changed bacterial peptides. According to this standard, comparing with the initial blank samples at day 0, a total of 953 differential bacterial peptide segments were obtained, of which, 513 peptide segments (53.83%) were up-regulated and 440 (46.16%) were down-regulated. The distribution of these up-regulated and down regulated bacterial peptide segments among dietary treatment groups was shown in Fig. 5.

**Fig. 5.**
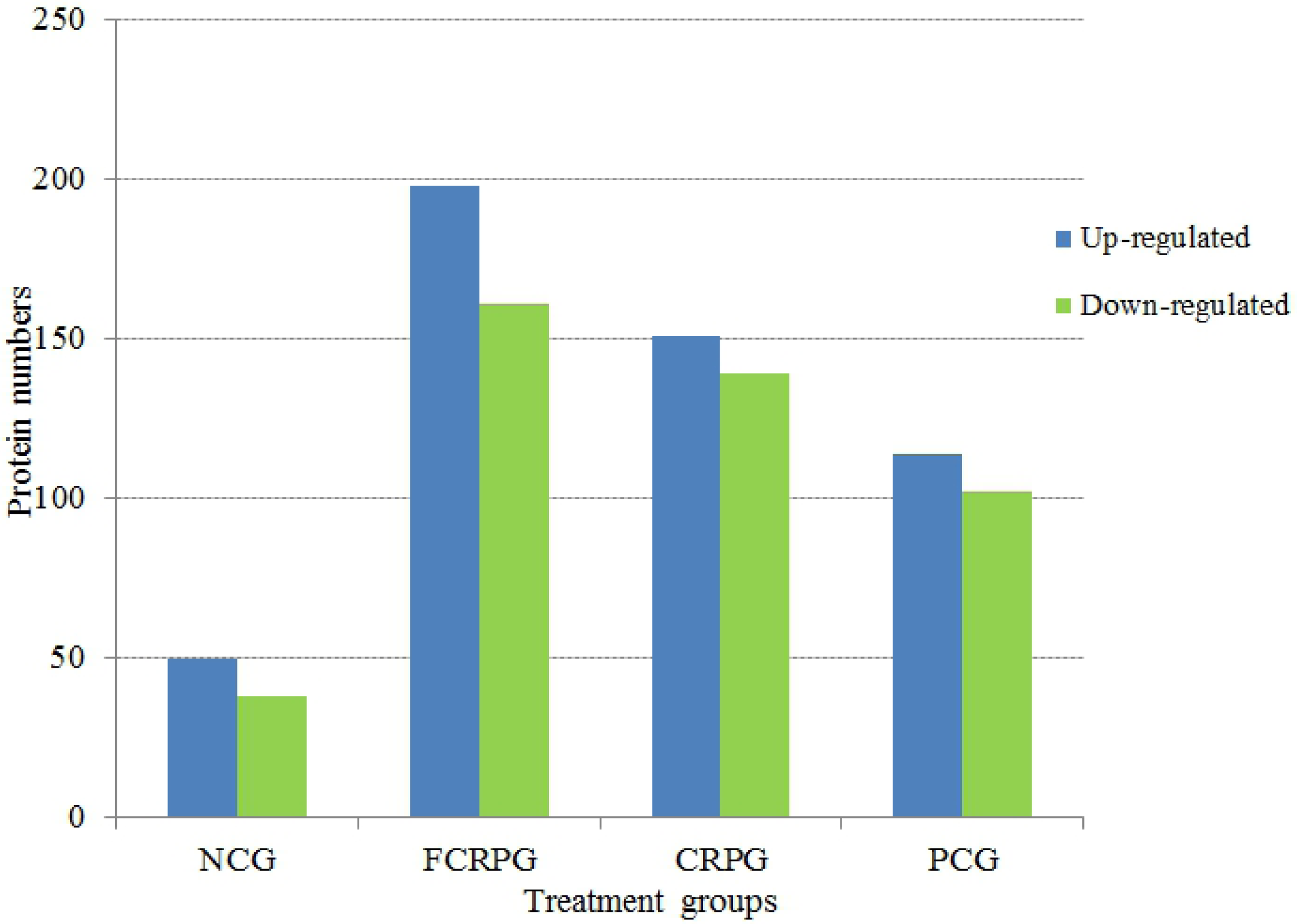
Distribution of bacterial peptide segments showing the number of up-regulated and down-regulated proteins among four dietary treatment groups.

### Analysis of the function of COGs of fecal bacteria proteins

As shown in Fig. 6, a total of 230 bacterial proteins in four treatment groups (NCG, FCRPG, CRPG and PCG) were assigned into 20 COGs. Among them, four predominant abundant functional categories are in turn as follows: B. Carbohydrate transport and metabolism, H. Energy production and conversion; I. Function unknown; T. Translation, ribosomal structure and biogenesis. In four treatment groups, FCRPG has the largest number of proteins that associated with the function of carbohydrate transport and metabolism, while CRPG contained the highest amount of proteins related to secondary metabolites biosynthesis, transport and catabolism.

**Fig. 6.**
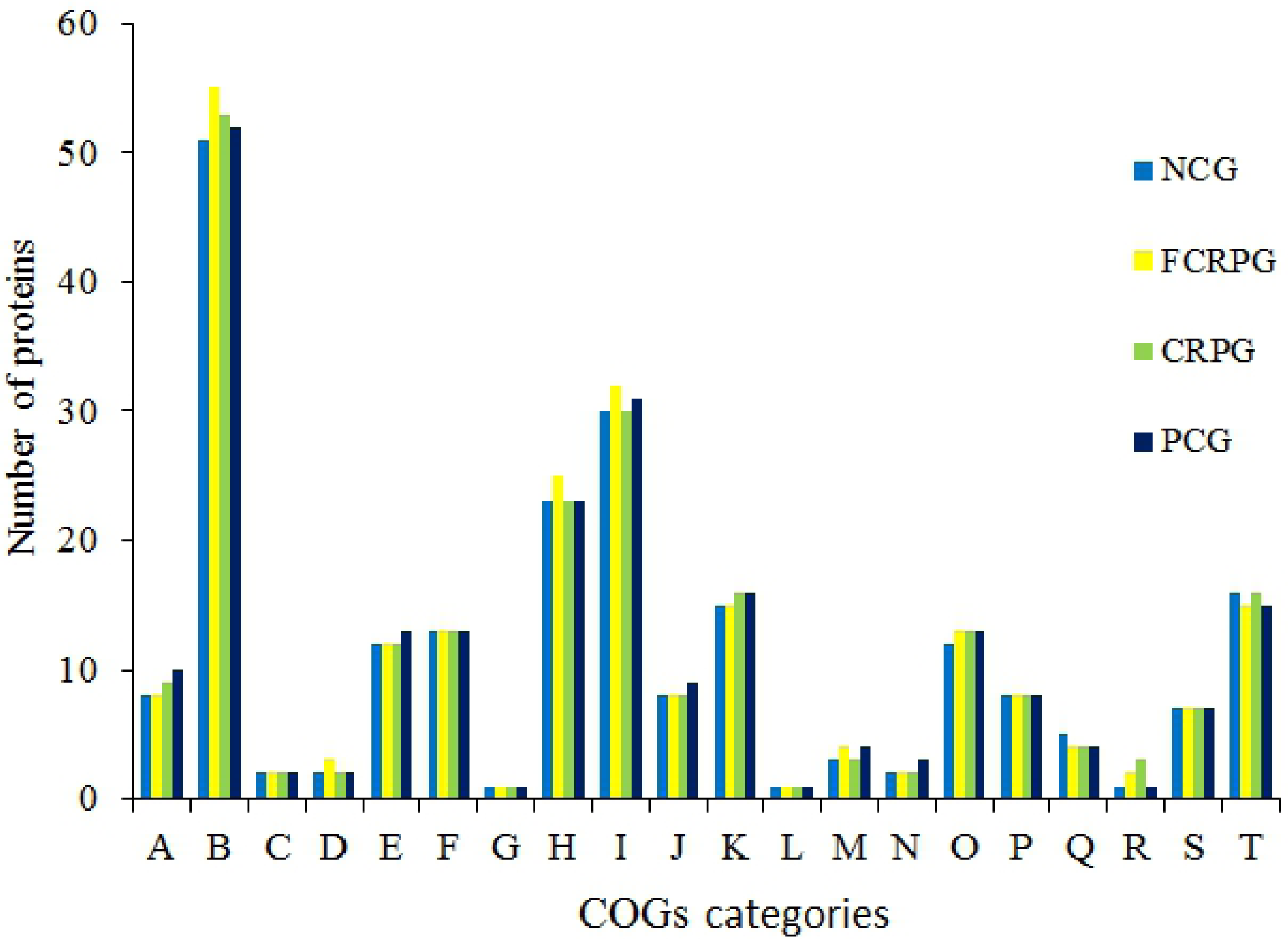
Analyses of COGs categories showing the functional differences of fecal bacterial proteins among dietary treatment groups. Note: A. Amino acid transport and metabolism; B. Carbohydrate transport and metabolism; C. Cell cycle control, cell division, chromosome partitioning; D. Cell motility; E. Cell wall/membrane/ envelopebiogenesis; F. Coenzyme transport and metabolism; G. Defense mechanisms; H. Energy production and conversion; I. Function unknow; J. General function prediction only; K. Inorganic ion transport and metabolism; L. Intracellular trafficking, secretion, and vesicular transport; M. Lipid transport and metabolism; N. Nucleotide transport and metabolism; O. Post-translational modification, protein turnover, chaperones; P. Replication, recombination and repair; Q. Signal transduction mechanism; R. Secondary metabolites biosynthesis, transport and catabolism; S. Transcription; T. Translation, ribosomal structure and biogenesis.

### KEGG pathway analysis of fecal bacterial proteins

Pathway-related annotation and analysis promote the in-depth understanding of biological function of identified proteins [2, 36]. In present study, the commericial KEGG database (https://www.kegg.jp/) was applied to annotate all of 230 differential bacterial proteins to extract functional informations. In accordance with the number of proteins involved in each pathway, 16 relatively major metabolic pathways were identified (Fig. 7). These pathways are mainly asscoiated with fatty acid metabolism, sugar metabolism, carbon metabolism and amino acid metabolism, etc. KEGG functional analysis showed that, among four dietary treatment groups, there have same number of functional proteins respectively in DNA replication and purine metabolism, while different number of proteins were observed in terms of fatty acid metabolism and butyrate metabolism. Specifically, FCRPG contains the relative maximum number of proteins pertaining to fatty acid metabolism and gastric acid secretion, while CRPG has the highest amount of proteins in flavonoid and flavonal biosynthesis.

**Fig. 7.**
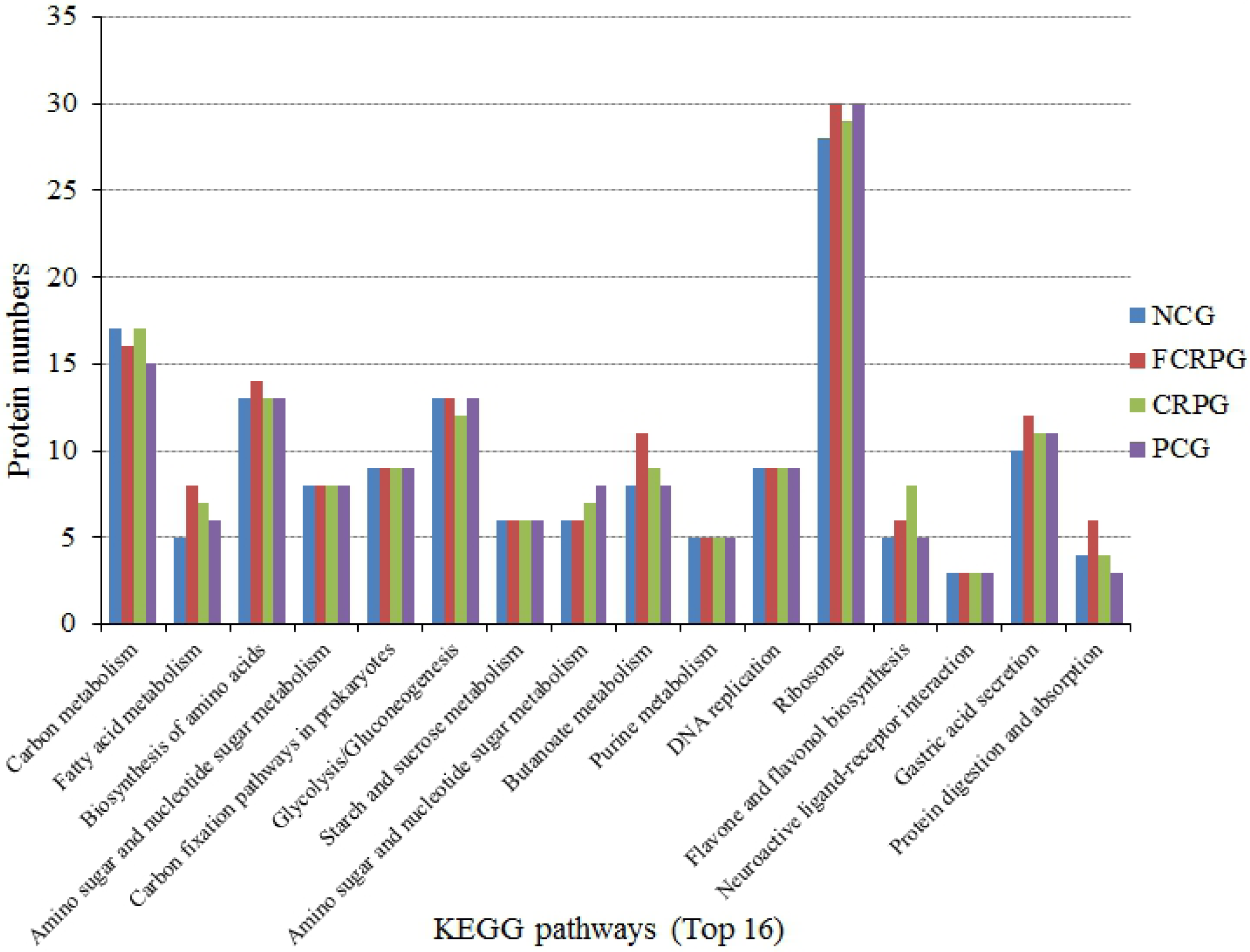
KEGG pathway analyses showing the differential fecal bacterial proteins. Top 16 metabolic pathways of KEGG were selected according to the number of proteins in corresponding pathway.

## Discussion

Understanding the critical role of the GI microbiota in host-microbe interactions during weaning transition in commercial piglets is of a great importance for reducing the risk of post-weaning infections (weanling piglets’ diarrhea) and promoting health and growth development. Previously published research suggested that gut microbiome, as a magical bioreactor, provides essential nutrient components like biotin and vitamin K, digests complex dietary fibres, as well as produces active metabolite SCFAs that nourish the gut epithelia [38]. However, little is still known to researchers about alimentary microbiome which has been recognized as “microbial dark matter” before. To our knowledge, the development of inflammatory GI diseases are obviously linked to alternations of digestive tract microbial communities [41]. On the other hand, the diversity and development of GI tract microbiome of piglets during weaning are basically frequently resulted from the changes of diets and physiological states [42]. Therefore, intestinal microorganisms play a role as a junction hub from diet intake to host physiological states (health or disease). Recently, technological advances of various omics data acquired from genomics, transcriptomics, metabolomics and proteomics have broaden the detailed investigations of intestinal microbiota for animals [3, 4]. These technologies hold great promise to provide robust strategies for scientists to elucidate the molecular mechanisms involved in host-microbial interactions in the complex feces ecosystem of weaned piglets.

### Correlation between functions and structures of fecal microbiome

In animals, microorganisms resided in alimentary canal are involving in food digestion, nutrient absorption, microbial fermentation and feces excretion. Therefore, the symbiotic microbita composition and its metabolic constituents have been considered as important factors for the characterization of GI tract health. In particularly, the first year of life is significant important in shaping and establishing the gut microbiota [45] which influencing the future life of animals. In the piglets’ feces, Firmicutes and Bacteroidetes are extensively regarded as the dominant phyla, followed by Proteobacteria and Actinobacteria, however, the abundance and composition of bacterial phyla was fluctuated and impacted by several determinants [46].

The diet compositions, to a certain degree, manipulate the composition and function of enteric microbiome. Specific microbial species/genus tend to be responsible for degradation of specific dietary components [47]. For example, two gut Bacteroides, B. thetaiotaomicron and B. ovatus, are capable to degradate almost all the main plant and host polysaccharides, including other microbial indigestible rhamnogalacturonan II [48]. Simultaneously, there also exists synergistic effects among the different bacteria genus. For example, Escherichia coli create an anaerobic circumstance beneficial to the colonized establishment of other bacterial species such as Bacteroides, Lactobacillus and Clostridium [49]. According to our data, FCRPG have a relative highest abundance of Acidobacterium in feces, followed by CRPG, compared with NCG and PCG, which suggest that dietary inclusion of FCRP or CRP could increase the contents of Acidobacterium to optimizate the intracavity environment of GI tract. In addition, the function category analysis of COGs show that FCRPG present a relative highest percentage value in the category of carbohydrate transport and metabolism, which hint that FCRP probably have an enhanced capability of strengthening the sugar metabolism.

### Bacteria-produced metabolite SCFAs

SCFAs play an important role on improving intestinal health in pigs [50]. It has been reported that SCFAs (e.g., acetate, propionate and butyrate) as end molecules derived from metabolism of soluble fibers by commensal microbiota exert multiple influences on gut morphology and function [51].

Butyrate that mainly produced by clusters IV and XIVa of Clostridia via the butyryl-CoA/acetate-CoA transferase enzyme, is preferentially used as an effective energy source for colonic epithelial cells, and play role in maintaining the intestinal homeostasis through multiple mechanism [52]. A remarkable decrease in butyrate production of gut microbiota is usually associated with certain functional disorders. For example, Fabry disease is often attributed to the accumulation of globotriaosylsphingosine (lyso-Gb3) which alters the formation of SCFAs, resulting in a significant reduction in butyrate concentration [53]. The present study recorded the significant highest fecal butyrate concentration (*P* < 0.05) in FCRPG, and the second highest value (*P* < 0.05) was observed in CRPG, which means that dietary supplemented with FCRP or CRP can evidently increase epithelial energy supply and maintain normal celluar proliferation and differentiation.

Propionate is mainly metabolized in the liver through gluconeogenesis, and as a potent growth stimulator for certain bacteria (e.g., Bifidobacterium). In addition, propionate has been shown to be able to bind to several receptors (e.g., GPR41 and GPR43) [51]. A new study found that propionic can serve as a powerful medicinal immunomodulatory supplement for multiple sclerosis patients in virtue of its normalization of Treg cell mitochondrial function and morphology in multiple sclerosis [54].

Acetate is widely produced by various bacterial taxas in mammalian gut. Both endogenous and indogenous, acetate has a beneficial effect on the function of host epithelial cells [55]. Previously, Lawhon et al. (2003) discovered that acetate can function as an inducer of invasion gene expression via a BarA/SirA-independent pathway [56, 57]. Recent study has pointed out that acetate fight against RSV-induced disorders via a mechanism of participating in activation of the membrane receptor GPR43 [58].

Associated metabolics study suggested that a reduced production of SCFAs has seen in antibiotic-treated mice and humans [59]. In our study, a significant decline of SCFAs content (*P* < 0.05) was observed in antibiotic intervention group (PCG) compared to FCRPG and CRPG, while no significant difference (*P* > 0.05) was observed when contrast to NCG.

### Bacteria-produced metabolite flavonoids

Flavonoid metabolism has been increasingly recognized for their crucial role in gut micro-bacteria. The composition of gut microbiota might be an important factor that affecting absorption of flavonoid-derived compounds by the animal host [60]. Four bacterial phyla (Actinobacteria, Firmicutes, Bacteroidetes, and Proteobacteria) are implicated in the bio-conversion of flavonoids, of which majority of bacterial species are capable of carrying out the O-deglycosylation of flavonoids for flavonoid transformation [61]. On the other hand, study in mouse models suggested that exogenous flavonoids intake can counteract the increased capacity of the microbiome to metabolize flavonoids, and do not hinder the anti-obesity functions of flavonoids in vivo [62]. Therefore, nutrition strategy is so important for modulating the intestinal microbial structure and thus result in regulating the level of metabolite flavonoids to maintain the health of animals. In present investigation, KEGG pathways analyses showed that CRPG has a highest number of bacterial proteins associated with flavone and flavonol biosynthesis, followed by FCRPG, compared to NCG and PCG.

Taken together, FCRP or CRP not only directly modulate the physicochemical characteristics of the digesta which interacting with intestinal mucosa, but also as the new growth substrates for particular microbacterial species so as to be beneficial for the health of weaned piglets.

## Conclusion

Essentially, CRP used in this work as a medical herb to exert its efficacy. FCRP can be considered as a kind of combination of prebiotics and probiotics. This study indicate that dietary inclusion of product of Citrus reticulata “Chachi” (FCRP or CRP) not only drives the alterations of fecal microbes populations but also modulates the microbial metabolic profiles to ameliorate the intestinal immune functions, which contributing to decode the biochemical mechanism of FCRP or CRP that beneficial to intestinal health of weaned piglets.

The utilizations of CRP-based herb compound recipe (e.g., Chenpisan) for promoting health and preventing diseases have been commercialized in swine production of China. Viewed from our observations, FCRP are capable to stimulate the appetite, while CRP can improve the carcass quality meat quality. Our results show that both FCRP and CRP, acting as antibiotic alternatives, modulate the microflora composition, microbiome phylotypes, and fecal microbial metabolites in their respective modes of action. This study may provide fundamental theoretical basis for the applications of FCRP or CRP in modulating microbiota of weaned piglets to facilitate health. Moreover, more studies are needed to carry out on combining diversified traditional Chinese herbal medicine to enhance the biological nutrition effect of FCRP or CRP so as to benefitial to piglets nursing. Finally, it should be noted that further directions to dissect the molecular mechanism in which how these medicinal plant products interaction with intestinal micro-ecosystem of piglets is of great significance to the national swine production.

## Acknowledgments

This investigation was supported in part by grants from the National Key Research and Development Plan (2016YFD0500600), National Key R & D Program Projects (2017YFF0104904), and Guangdong Provincial Science and Technology Plan Project (2017B020207004). Additionally, we thank Ph.D. student Xusheng Li for offering assistances in drawing figures.

